# Phylogenomics of the major tropical plant family Annonaceae using targeted enrichment of nuclear genes

**DOI:** 10.1101/440925

**Authors:** Thomas L.P. Couvreur, Andrew J. Helmstetter, Erik J.M. Koenen, Kevin Bethune, Rita D. Brandão, Stefan Little, Hervé Sauquet, Roy H.J. Erkens

## Abstract

Targeted enrichment and sequencing of hundreds of nuclear loci for phylogenetic reconstruction is becoming an important tool for plant systematics and evolution. Annonaceae is a major pantropical plant family with 109 genera and ca. 2450 species, occurring across all major and minor tropical forests of the world. Baits were designed by sequencing the transcriptomes of five species from two of the largest Annonaceae subfamilies. Orthologous loci were identified. The resulting baiting kit was used to reconstruct phylogenetic relationships at two different levels using concatenated and gene tree approaches: a family wide Annonaceae analysis sampling 65 genera and a species level analysis of tribe Piptostigmateae sampling 29 species with multiple individuals per species. DNA extraction was undertaken mainly on silicagel dried leaves, with two samples from herbarium dried leaves. Our kit targets 469 exons (364 653 bp of sequence data), successfully capturing sequences from across Annonaceae. Silicagel dried and herbarium DNA worked eaually well. We present for the first time a nuclear gene-based phylogenetic tree at the generic level based on 317 supercontigs. Results mainly confirm previous chloroplast based studies. However, several new relationships are found and discussed. We show significant differences in branch lengths between the two large subfamilies Annonoideae and Malmeoideae. A new tribe, Annickieae, is erected containing a single African genus *Annickia*. We also reconstructed a well resolved species-level phylogenetic tree of the Piptostigmteae tribe. Our baiting kit is useful for reconstructing well supported phylogenetic relationships within Annonaceae at different taxonomic levels. The nuclear genome is mainly concordant with plastome information with a few exceptions. Moreover, we find that substitution rate heterogeneity between the two subfamilies is also found within the nuclear compartment, and not just plastomes and ribosomal DNA as previously shown. Our results have implications for understanding the biogeography, molecular dating and evolution of Annonaceae.

## Introduction

Targeted enrichment followed by high throughput sequencing of hundreds or even thousands of loci for phylogenetic reconstruction is becoming a golden standard in plant evolutionary biology (Cronn et al., 2012; Mariac et al., 2014; Barrett et al., 2016; Xi et al., 2013). Targeted capture is a genome reduction approach whereby selected regions of the genome are “captured” or hybridized in solution using site-specific baits (also called probes). Once the baits hybridize to the targeted regions, the rest of the genome is discarded, and only the regions of interest are sequenced (Grover et al., 2012) increasing the read depth of the regions of interest. This is in contrast to earlier approaches where the focus lay on recovering complete genomes (e.g. Staats et al., 2013). Multiplexing approaches based on individual barcodes enable fast and cost effective sequencing of multilocus sequence data for phylogenetic or phylogeographic inference (McCormack et al., 2013). Targeted capture of sequence data, either nuclear or plastid, has been used to reconstruct phylogenetic relationships at several taxonomical scales from within angiosperms (Stull et al., 2013; Johnson et al., 2018) to within families (Barrett et al., 2016; Mandel et al., 2014), or to increase phylogenetic resolution in species-rich clades (Nicholls et al., 2015) or at infra-specific levels (Faye et al., 2016; Zellmer et al., 2012). The difficult step concerns the identification of the targeted regions and the design of the baits. This relies on genome-wide information such as transcriptomes or full genomes to identify low copy orthologous genes useful for phylogenetic inference. These data are not always available for non-model groups or clades, especially for tropical lineages. However, increased availability of transcriptomes across angiosperms (Wickett et al., 2014) and lowered sequencing costs allow access to more genomic resources useful in non-model groups.

Annonaceae is a pantropical family of trees, shrubs and lianas (Chatrou et al., 2012a) and is among the most species-rich families of tropical rain forest lineages (Chatrou et al., 2012a). To date Annonaceae contain 109 genera and around 2430 species (Chatrou et al., 2012b; Guo et al., 2017b; Xue et al., 2018). Phylogenetic analyses of the family started in the 1990’s with a morphology-based phylogenetic tree published by Doyle and Le Thomas (1994) providing a first cladistic understanding of relationships within the family. In the molecular era, chloroplast sequence data have been the main markers for Annonaceae family and species level phylogenetic reconstructions (Doyle et al., 2000; Mols et al., 2004; Richardson et al., 2004; Pirie et al., 2006; Su et al., 2008; Couvreur et al., 2011a; Chatrou et al., 2012b; Guo et al., 2017a,b; Thomas et al., 2017). Most of these studies, however, used a relatively small number of markers (but see Hoekstra et al. (2017); Lopes et al. (2018); Guo et al. (2018) for notable exceptions). For instance, the latest classification update of Annonaceae is based on eight chloroplast markers (Chatrou et al., 2012b). Although this led to an overall well supported generic phylogenetic tree, several relationships remained weakly supported. In addition, species level phylogenies of different Annonaceae genera are, in general, moderately supported based on plastid markers alone (e.g. Pirie et al., 2006; Erkens et al., 2007; Thomas et al., 2017; Couvreur et al., 2011b).

Annonaceae is subdivided into four subfamilies and 15 tribes (Chatrou et al., 2012b; Guo et al., 2017b). Subfamilies Annonoideae and Malmeoideae contain over 90% of all species in the family (1515 and 783, respectively (Guo et al., 2017b) when compared to the other two smaller subfamilies Anaxagoreoideae and Ambavioideae. Based on plastid phylogenetic analyses, Annonoideae and Malmeoideae showed contrasting overall branch lengths, initially leading to them to be known as “long branch” and “short branch” clades, respectively (Richardson et al., 2004). Recently, it has been shown that these phylogenetic branch differences are linked to different substitution rates within the chloroplast and ribosome (Hoekstra et al., 2017).

To date, no nuclear phylogenetic Annonaceae tree has been generated, mainly because of difficulties in amplifying nuclear genes using standard Sanger approaches (Pirie et al., 2005). In this study, we aimed to generate for the first time a nuclear phylogenetic tree of Annonaceae at both generic and species level using target sequence capture and next generation sequencing. We designed and tested a baiting kit useful at several taxonomic levels with the ultimate aim of reconstructing the Annonaceae Tree of Life, including all species in a well-resolved phylogenetic tree based on a wide coverage of the nuclear genome. Specifically, we wanted to answer several questions: Can we generate a baiting kit useful for Annonaceae wide phylogenetic analyses? Are plastid and nuclear phylogenetic relationships concordant in Annonaceae? Do we detect differences in branch lengths based on nuclear data between Annonoideae and Malmeoideae?

## Material and Methods

### Nuclear bait design

#### Sampling

Nuclear baits were designed based on the analysis of transcriptomes sequenced from five species of Annonaceae, two from Malmeoideae and three from Annonoideae (Table 1). For four species, leaf material was collected at the Utrecht Botanical Garden (The Netherlands) and stored in RNAlater storage solution (Sigma-Aldrich, St. Louis, USA). Flowers of *Marsypopetalum littorale* (DIV030) were collected with forceps at the time of anthesis on July 11 2013 at the Botanic Gardens, Vienna (Austria). Flowers were placed in Falcon tubes and immediately submerged in liquid nitrogen, then stored at −80°C.

**Table 1.**
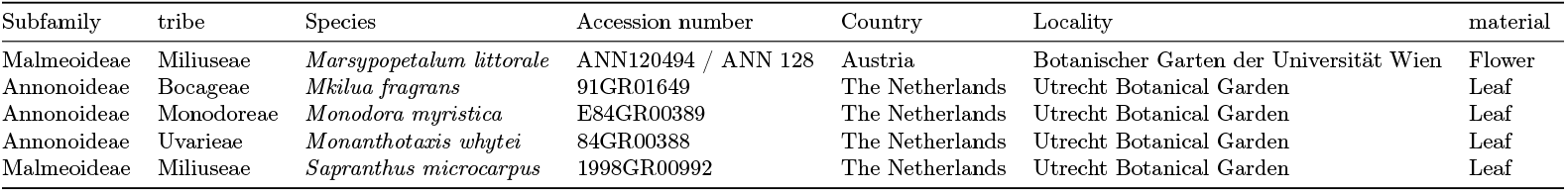
Species used to sequence transcriptomes and design the nuclear bating kit

#### RNA extraction

Each leaf piece was homogenized using liquid nitrogen. RNA isolation was conducted using a CTAB approach and a two-step chloroform:isoamylalcohol (24:1) extraction. RNA was finally eluted in RNase-free water and treated with DNase I (Qiagen, Hilden, Germany). RNA extraction from floral material was based on 100 mg of material and followed modified protocols (Kolosova et al., 2004; Dal Cin et al., 2005). RNA pellets were suspended in RNAse free water.

#### RNA quantification, library preparation and sequencing

RNA was quantified in a Qubit 2.0 instrument (Invitrogen, Carlsbad CA, USA) using the Quant-iT RNA assay kit (Invitrogen, Carlsbad CA, USA) according to the manufacturer’s protocol. A total of 4 g of RNA was used for library preparation using the TruSeq^®^ Stranded mRNA Library prep kit (Illumina, San Diego CA, USA) following the manufacturer’s protocol for low throughput samples (Part 15031047 Rev. E). An alternate fragmentation protocol was followed to obtain insert sizes of 200 bp as described in the appendix of the manufacturer’s protocol. The obtained libraries were quantified using a Qubit 2.0 instrument (Invitrogen, Carlsbad CA, USA) and Qubit dsDNA HS assay kit (Invitrogen, Carlsbad CA, USA). Samples were pooled to a final concentration of 4 nM and sequenced on an Illumina HiSeq 2000 sequencer (Illumina, San Diego CA, USA) using 2 × 100 bp chemistry.

#### Bait design

Reads generated from the transcriptome of each species were assembled *de novo* using Trinity (Grabherr et al., 2011). Using the resulting contigs, orthologous loci were selected following these steps. First, candidate Open Reading Frames (ORF) were extracted and translated using TransDecoder **https://github.com/TransDecoder** resulting in a number of exons (start and stop codons present). The selected exons were then clustered together based on protein similarity using CD-HIT (Li and Godzik, 2006; Fu et al., 2012). Finally, we used SelfBLAST to discard exons with multiple hits in the same species. Using this set of exons we applied a 5-way reciprocal best hit (RBH) comparison in order to identify those shared between 3, 4 and 5 species. This step was undertaken twice. Once with *Monodora myristica* (Annonoideae) and once with *Marsypopetalum littorale* (Malmeoideae) as focal taxa against which other taxa were compared. This allowed us to identify set of loci common within each subfamily and amongst Annonaceae. Shared loci were then aligned using MAFFT (Katoh and Standley, 2013), subsequent alignments were trimmed using BMGE (Criscuolo and Gribaldo, 2010), and phylogenetic relationships were constructed using RAxML (Stamatakis, 2014). We only selected contigs that reconstructed phylogenetic relationships congruent with known Annonaceae relationships between these five taxa (Chatrou et al., 2012b; Guo et al., 2017b). Indeed, our sampling of transcriptomes comes from phylogenically distant Annonaceae species from two major subfamilies. We do not expect that these relationships based on plastid markers be different at the nuclear level.

We further filtered these identified low copy orthologous sequences by selecting exons that were longer than 300 bp. Then, we excluded homologous sequences where the percentage identity between different pairs was different from what we expect from known evolutionary relationships of those taxa. Following the phylogenetic tree of Annonaceae (Chatrou et al., 2012b; Guo et al., 2017b) we expect *((Mkilua,(Monodora, Monanthotaxis)),(Marsypopetalum, Sapranthus)).* Regions with a percentage identity of two phylogenetically closest species >0.75 and two furthest species >0.7 were selected. For sequences longer than 1500 bp we were more stringent (identity of closest species >80). Finally, we excluded regions with a high variability of identity percentage (variance >130).

One particularity is that we kept redundant exon sequences between species when possible. Thus if a same exon region was present in four different species, baits were generated for all four “variants” of that exon. This was done to increase capture efficiency across the whole family for that exon. This is similar in concept to a degenerated primer pair for example.

Final baiting sets were synthesized at Arbor Biosciences (MYbaits **http://www.arborbiosci.com/**) to 120 bp long, with 3X tiling and with a 40 bp spacing between the start of neighboring baits.

### Phylogenetic analyses of Annonaceae

#### Taxon sampling

In order to validate our nuclear-baiting kit for family-wide Annonaceae phylogenetics, we undertook a two-level analysis. First, we aimed to reconstruct, for the first time, a generic level nuclear phylogenetic tree of Annonaceae (referred to as “Annonaceae” analyses). We sampled 65 genera (out of 109) from all four subfamilies (see supplementary information Table 1) representing a total of 11 tribes out of the 15 currently recognized tribes (Chatrou et al., 2012b; Guo et al., 2017b). From one to 12 species were sampled per tribe. All genera were represented by one species, except for the pantropical genus *Xylopia* (two species included: one from South America, one from Africa) and the paraphyletic genus *Friesodielsia* (Guo et al., 2017a) for which we sampled one African species and one South-East Asian species.

Second, to assess its usefulness for reconstructing species-level relationships, we undertook a species-level sampling of the African tribe Piptostigmateae (referred to as “Piptostigmateae” analyses). This tribe comprises seven genera (Ghogue et al., 2017) most of them recently revised (Versteegh and Sosef, 2007; Couvreur et al., 2009, 2015; Marshall et al., 2016; Ghogue et al., 2017). We sampled a total of 29 out of 39 species covering all seven genera (see supplementary information Table 1). One to five individuals were sampled per species, plus a number of outgroup taxa, leading to a total of 83 individuals included in our analyses. All DNA was extracted from silicagel dried leaves, except for two samples, where DNA was extracted from air dried herbarium samples collected in 2000.

#### DNA extraction, library preparation, in-solution hybridization and sequencing

DNA was extracted from silica gel dried leaves following the MATAB and chloroform separation methods of Mariac et al. (2014) and Scarcelli et al. (2006). The full protocol is provided in supplementary information Protocol 1. Illumina libraries were constructed following a modified protocol of Rohland and Reich (2012) using 6-bp barcodes and Illumina indexes to allow for multiplexing at different levels. Extra steps were added to the Rohland and Reich (2012) protocol to allow for amplification and in-solution hybridization as follows: total DNA for each individual was sheared using a Bioruptor Pico (Diagenode, Liége, Belgium) to a mean target size of 500 bp. DNA was then repaired, ligated and nick filled-in before an 8 to 11 cycle prehybridization PCR was performed. After clean-up and quantification, libraries were bulked, mixed with biotin-labelled baits and hybridized to the targeted regions using the bait kit designed above. The hybridized biotin-labelled baits were then immobilized using streptavidin-coated magnetic beads. A magnetic field was applied and supernatant containing unbounded DNA was discarded. Enriched DNA fragments were then eluted from the beads and amplified in a 14 to 16 cycle real-time PCR to complete adapters and generate final libraries. Libraries were sequenced on an Illumina HiSeq v3 platform pair end and length of 150 bp (Illumina, SAn Diego CA, USA) at CIRAD facilities (Montpellier, France) with around 18 pmol of the capture-amplified DNA libraries deposited on the flowcell. The full step-by-step protocol is provided in supplementary information Protocol 2.

### Bioinformatics

Demultiplexing with a 0-mistmatch threshold was undertaken using the demultadapt script (**https://github.com/Maillol/demultadapt**). Adapters were removed using cutadapt 1.2.1 (Martin, 2011) with the default parameters. Reads were filtered according to their length (>35 bp) and quality mean values (Q > 30) using a custom script (**https://github.com/SouthGreenPlatform/arcad-hts/blob/master/scripts/arcad_hts_2_Filter_Fastq_On_Mean_Quality.pl**). Forward and reverse sequences were paired according to their name in the fastq files using a comparison script, adapted from TOGGLe (Tranchant-Dubreuil et al., 2018). A terminal trimming of 6 bp was performed on reverse sequences to ensure removal of barcodes in case of sequences shorter than 150 bp using the fastx trimmer script which is part of the fastx toolkit (**https://github.com/agordon/fastx_toolkit**). A final sorting by sequence identifier was done on the stq file before further analyses.

### Contig assembly and multi-sequence alignment

We used the pipeline HybPiper (v1.2) (Johnson et al., 2016) to process our cleaned data. Briefly, reads were mapped to targets using BWA (v0.7.12) (Li and Durbin, 2009) and those reads that were successfully mapped were assembled into contigs using SPAdes (v3.11.1) (Bankevich et al., 2012). Exonerate (Slater and Birney, 2005) was then used to align the assembled contigs to their associated target sequence. Additionally, if contigs were slightly overlapping they were combined into ‘supercontigs’ which contained both target and off-target sequence data. Exonerate was run a second time so that introns could be more accurately identified. We aligned each set of supercontigs using MAFFT (v7.305) (Katoh and Standley, 2013) with the “-auto” option and cleaned these alignments with GBLOCKS (v0.91b) (Castresana, 2000) using the default parameters and all allowed gap positions. The number of parsimony informative sites were calculated for each supercontig alignment using the “pis” function in the ape R package (Paradis et al., 2004).

### Paralog identification

HybPiper flags potential paralogs when multiple contigs are discovered mapping well to a single reference sequence. The program uses coverage and identity to a reference to choose a ‘main’ sequence and denotes the others as potential paralogs. We took flagged loci and constructed gene trees using RAxML (v8.2.9) (Stamatakis, 2014). We examined each tree to determine whether putative paralogs formed a species clade, in which case these were unlikely true paralogs and we could continue with the main sequence selected by HybPiper. If the ‘main’ and alternative sequences formed separate clades they were likely true paralogs so we removed the entire locus from the downstream analyses.

### Phylogenetic inference

We inferred trees using a generic-level dataset of Annonaceae and a species-level dataset of the tribe Piptostigmateae (see Supplementary Table 1). For each dataset we identified those exons that had 75% of their length reconstructed in 75% of individuals. We then took the corresponding supercontigs (i.e. targeted exon regions plus surrounding off-target captured sequence data) for phylogenetic inference. We initially tried tree inference using exons only but support was generally lower than trees based on supercontigs and the topology was similar so we decided to proceed using supercontigs (results not shown). All trees were reconstructed unrooted (as is mandatory in ASTRAL) and then rooted post-inference. The Annonaceae tree was rooted on *Anaxagorea crassipetala* and the Piptostigmateae tree was rooted using *Annona glabra.*

### Coalescent approach

Individual gene trees were constructed using RAxML (v8.2.9) (Stamatakis, 2014) with a GTR+GAMMA model and 100 bootstrap replicates. Branches with boostrap support <10 were collapsed using Newick Utilities program nw_ed (Junier and Zdobnov, 2010). This has been shown to improve the accuracy of inferred species trees (Zhang et al., 2017). We then ran ASTRAL-III (Zhang et al., 2017) using the default parameters and the gene trees that corresponded to our selected loci.

### Concatenation approach

After alignment, we filled missing individuals at each locus with an empty sequence and concatenated aligned loci using the pxcat function in the program phyx (Brown et al., 2017). A GTR+GAMMA model was defined using the partition file output by pxcat leading to a different substitution model for each alignment (Brown et al., 2017). A rapid bootstrap analysis in RAxML (v8.2.9) (Stamatakis et al., 2008; Stamatakis, 2014) with 100 replicates was performed followed by a thorough ML search on the original alignment.

## Results

### Baiting kit

The trinity analyses recovered a total of 297 193 raw contigs for *Marsypopetalum littorale,* 342 592 for *Sapranthus microcarpus,* 102 025 for *Monodora myristica,* 194497 for *Monanthotaxis whytei* and 164 635 *Mkilua fragrans*. After the orthologous low copy selection process we ended up with 80 317, 68 039, 68 039, 22 326, 33 794 contigs, respectively. Targeted exon regions varied from 300 bp to 6072 bp. A total of 813 exonic regions were finally selected after our filtering process. However, a number of these regions were present several times as different variant of the same exon: 236 were present twice, 31 were present three times, and 15 present four times. Thus, the baiting kit effectively targets 469 unique exonic regions with a total capture footprint of 364 653 bp. The final MyBaits Annonaceae baiting kit was made of 11 583 baits 120 bp long.

### Number of loci retrieved and variability

Overall, we recovered good average depth for all individuals sequenced (275; min: 45; max: 726) and on average we recovered over 90% of our target with a coverage great than 10× (Supplementary table 1).

For the Annonaceae analyses, we recovered 98 to 468 loci depending on the percentage of reconstructed targeted exon length for the same percentage of individuals (Table 2). We recovered at least a fraction of all but one of the targeted loci (468 of 469). A total of 331 loci were reconstructed for over 75% of their length and for over 75% of individuals and this ‘75/75’ subset was used for tree inference. The total length of these 331 loci was 545,610 bp.

For the Piptostigmateae analyses, we captured between 170 and 469 loci (Table 2). We captured at least some of all the targeted exons in this dataset. A total of 379 loci were retrieved under the 75/75 rule and used for the phylogenetic inference. The total length of these 379 loci was 766,373 bp.

We recovered substantial amounts of non-target sequence data per locus. The typical aligned length of supercontigs was more than twice that of exon alignments (Figure 1). On average we recovered 828 bp of off-target sequence data in our Annonaceae dataset and 1241 bp in our Piptostigmateae dataset to be used for downstream phylogenetic inference.

We calculated 75/75 loci for each Annonaceae subfamily and identified 179 loci common across each subfamily locus set. We found loci that were shared between all but three subfamily combinations, and loci that were specific to each subfamily except Ambavoideae (Figure 2A). The set of 75/75 loci used in the Piptostigmateae tree inference contained almost all of those used in the Annonaceae tree inference as well as 51 additional loci (Figure 2B).

**Figure 1.**
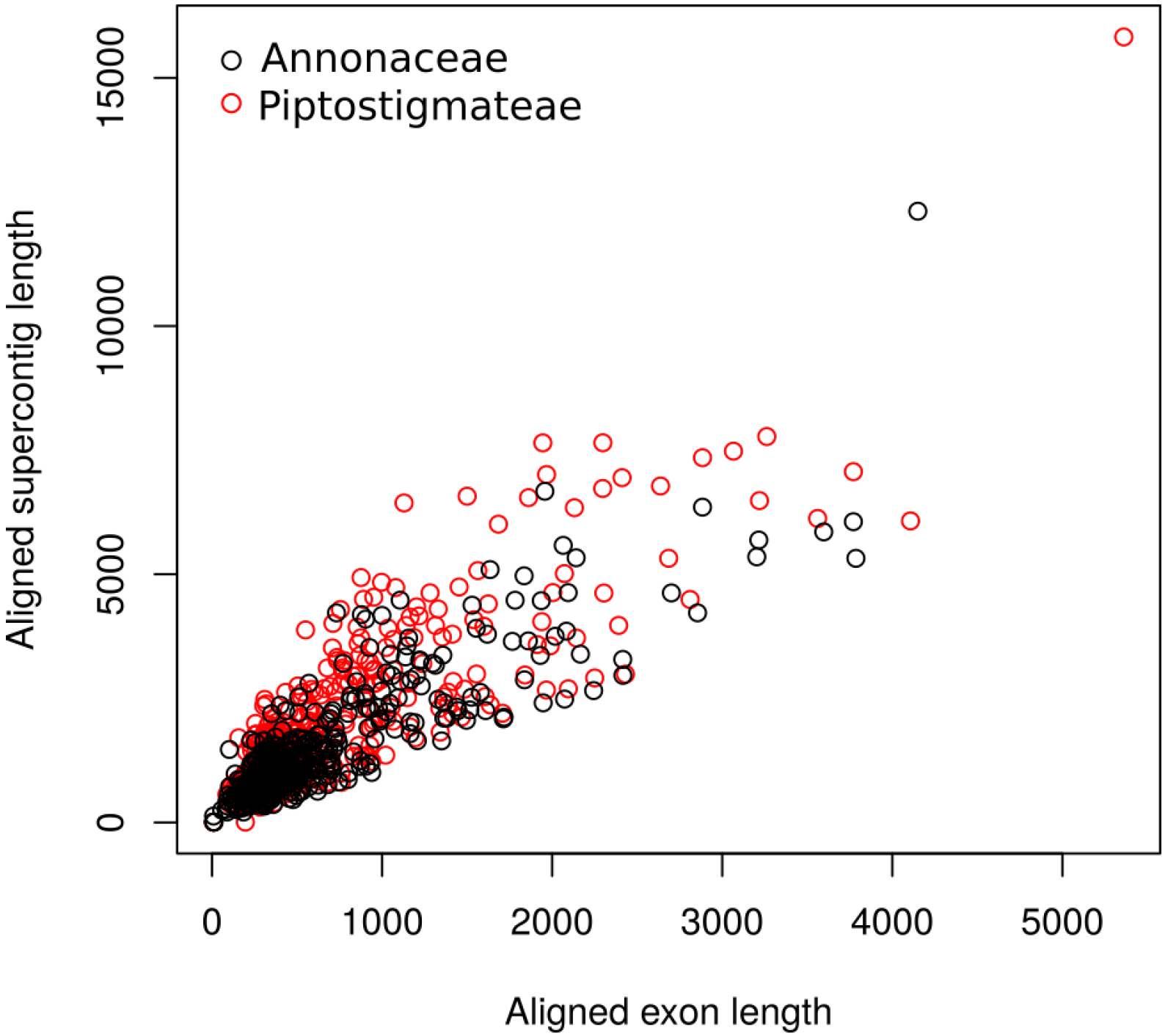
Scatterplot of targeted exon length against supercontig length (targeted exon plus off target data) at each locus. Lengths were calculated post-alignment. Black open circles: Annonaceae alignments; Red open circles: Piptostig-mateae alignments.

At the Annonaceae level, post alignment length of the 331 supercontigs varied from 407 to 12 314 bp long with a mean of 1648 bp (Table 3). For the Piptostigmateae matrix, the supercontigs varied from 492 to 15 824 bp long with a mean of 2 022 bp (Table 3). In both cases, the longer the supercontigs the more Parsimony Informative Sites were retrieved (Fig. 3C).

### Family-level analyses

We visually inspected loci flagged as putative paralogs by hybpiper to verify whether there was a clear phylogenetic distinction between main and alternative paralogs. This lead to 14 loci where paralogy seemed probable and these were removed from our 75/75 locus subset. After removal, the post alignment concatenated length of the remaining 317 loci was 533 322 bp. Mean supercontig length was 1648 bp long, with on average 861 parsimony informative sites and the longest supercontig was 12 314 bp long (3).

Support was generally high throughout the Annonaceae ASTRAL tree, with approximately 86% of branches possessing local posterior probabilities (LPP) of more than 90% (Figure 4 A). An estimated 95% of input quartet trees were in the species tree indicating a generally low level of gene tree conflict in the family. However, assessing quartet support at nodes (Figure 4 B) revealed gene tree conflict was high in the Miliuseae. The RAxML tree (Figure 5) was inferred with very high overall levels of bootstrap support - 88% of branches had 100 bootstrap support. The topologies produced by the ASTRAL and RAxML approaches were very similar. RAxML consistently conferred higher support in instances of gene tree conflict such as the placement of Miliuseae genera. One of the most conspicuous differences between approaches is the placement of the genus *Annickia.* This genus is inferred to be sister to the Malmeoideae in the ASTRAL tree while it is sister to the rest of the Piptostigmateae in the RAxML tree, albeit with low support. In addition, several genera are recovered in different positions than previously inferred using plastid markers, such as *Artabotrys* and *Sanrafaelia*.

**Table 2.** Variability of number of captured of loci using the Annonaceae bait kit. The values indicate the percentage of individuals for which we reconstructed the same percentage of the targeted exon region. Phylogenetic analyses were carried out using the 75/75 datasets

|  | > 0% | ≥ 25% | ≥ 50% | ≥ 75% | 95% |
| --- | --- | --- | --- | --- | --- |
| Annonaceae | 468 | 466 | 449 | 331 | 98 |
| Piptostigmateae | 469 | 466 | 453 | 379 | 170 |

**Figure 2.**
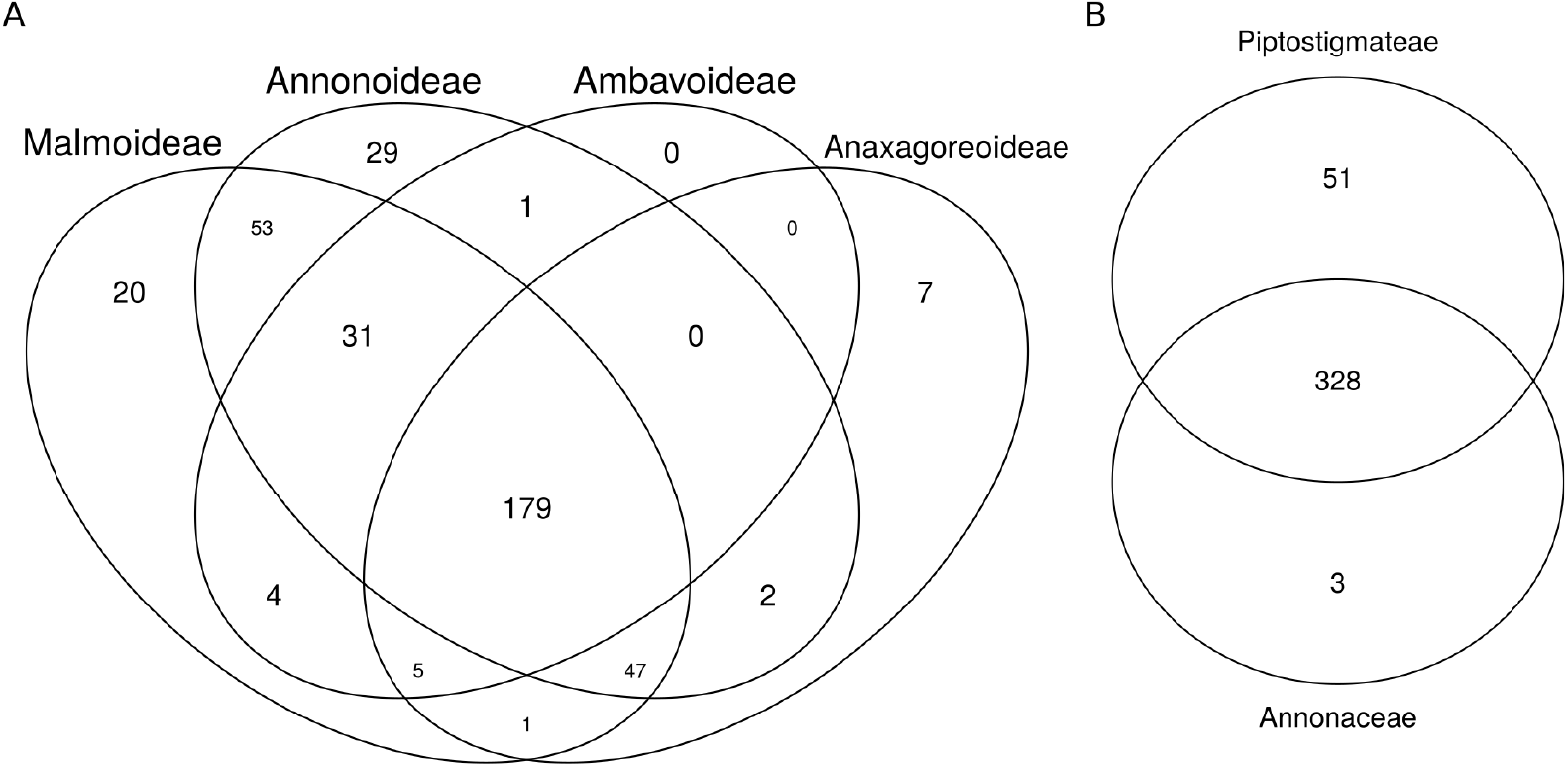
Venn diagrams of shared loci between major Annonaceae clades. A. Annonaceae diagram: Individuals from the Annonaceae analyses were separated based on subfamily groups. The number of loci sequenced for more than 75% of the targeted exon length and present in more than 75% of individuals was calculated for each subfamily. The diagram thus represents the number of shared phylogenetically useful loci within and among Annonaceae subfamilies. B. Piptostigmateae diagram. Reliable loci (calculated as above) used in Annonaceae and Piptostigmateae tree inference were compared to identify the number of overlapping and unique loci used at each level.

### Piptostigmateae analyses

After visual verification we identified 24 probable paralogs, 23 of which were found in our 75/75 set of loci and removed from downstream analyses. The remaining 356 loci contained a total of 743,325 bp of sequence data. Mean supercontig length was 2022 bp long, with on average 743 parsimony informative sites and the longest supercontig was 15824 bp long (3).

**Figure 3.**
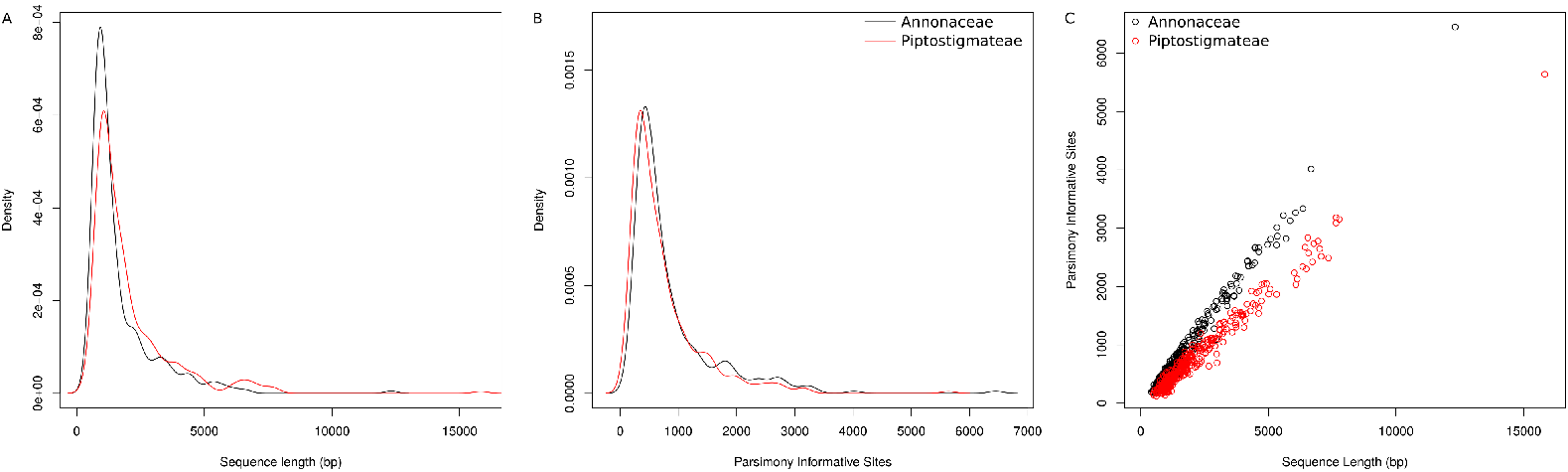
Information content of targeted sequences for the whole of the Annonaceae family (in black) and for the tribe Piptostigmateae (in red). Values provided are post alignment of supercontigs (targeted exon plus off target data). A: Density plot of supercontig length. B: Density plot of Parsimony Informative Sites. C: Scatter plot of supercontig length and Parsimony Informative Sites

**Table 3.** Length and parsimony informative sites statistics based on the aligned Annonaceae and Piptostigmateae matrices

|  |  | Supercontig | PIS |
| --- | --- | --- | --- |
| Annonaceae | Mean | 1 648.4 | 861.7 |
|  | SD | 1 325.7 | 746.6 |
|  | Min | 407 | 169 |
|  | Max | 12 314 | 6 450 |
| Piptostigmateae | Mean | 2 022.1 | 743.4 |
|  | SD | 1 621.7 | 637.9 |
|  | Min | 492 | 122 |
|  | Max | 15 824 | 5 639 |

We found similar high levels of node support in our Piptostigmateae trees when compared to the Annonaceae analyses, despite the reduced evolutionary distance between taxa (Figure 6 A). 85% of branches had LPP values greater than 0.9. Gene tree conflict was again low (91% of gene tree quartets were represented in the species tree) but slightly higher than in the Annonaceae tree. This conflict was principally found within species (Figure 6 B), for example *Piptostigma goslineanum* formed a well supported clade but branches within possessed low quartet support and LPP. A small number of nodes found deeper in the genera *Greenwayodendron* and *Piptostigmata* were also poorly supported (Figure 6 A). As in the Annonaceae analysis, we found increased support in all areas of the Piptostigmateae RAxML tree and the topologies generated by the two inference methods were similar. Likewise, RAxML tended to give higher levels of support in those areas of the tree where ASTRAL estimated low LPP. Like in the Annonaceae tree the placement of *Annickia* changed between approaches either sister to (RAxML) or diverging before (ASTRAL) the rest of the Piptostigmateae split from the rest of the Malmeoideae genera sampled.

**Figure 4.**
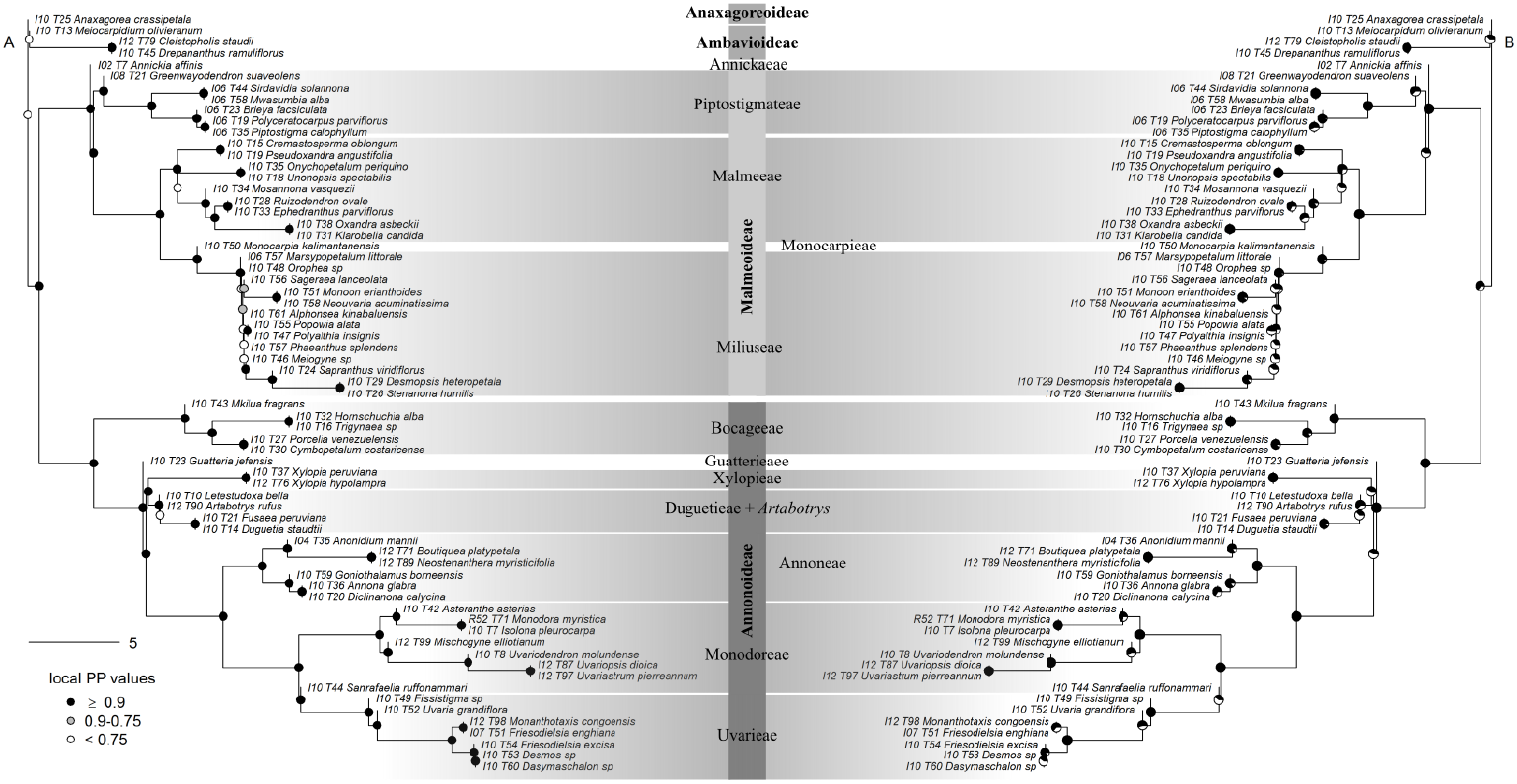
Generic-level tree of Annonaceae constructed using ASTRAL. A.Tree inference was based on 317 supercontigs (exons & introns). Depicted on nodes are the local posterior probability (LPP) values. B. Identical tree to A but with quartet support represented on nodes. Black portion of pie charts represents the percentage of quartets in gene trees agreeing with this branch.

## Discussion

### An Annonaceae-wide nuclear baiting kit

Generating “baiting kits” for targeted enrichment of hundreds of nuclear loci using Next Generation Sequencing is becoming a major component of plant phylogenetic studies (Nicholls et al., 2015; Cronn et al., 2012; Zellmer et al., 2012; Heyduk et al., 2016; Johnson et al., 2018; Kadlec et al., 2017). Here, we present a baiting kit that can be used to undertake phylogenetic studies across the large pantropical Annonaceae family at different taxonomic levels (e.g. Figures 5, 7). The Annonaceae kit enables potential capture of 469 unique exonic regions (Table 2). Although the kit was designed using comparative transcriptomic analyses of five species from two of the most diverse Annonaceae subfamilies (Annonoideae and Malmeoideae), we successfully captured shared loci within the two other smaller subfamilies (Ambavioideae and Anaxagoreoideae). Under our 75%/75% rule (see methods), 179 loci were common across all major subfamilies (Figure 2 A). However, few loci were recovered unique to Anaxagoroideae (7) and none for Ambavoideae (Figure 2 A). These results indicate that this baiting kit has a wide taxonomic breadth of utility, and will be useful to reconstruct phylogenetic relationships across the family. We show that our baiting kit is effective at recovering a similar number of loci when we focus on a specific tribe with a species level sampling. For the Piptostigmateae tribe, and before removal of potential paralogues, we recovered 379 loci under the 75/75 rule (Table 2). Thus, at a lower taxonomic level we recovered slightly more loci than at the family level (331 loci at 75/75). In addition, 87% of loci recovered for Piptostigmateae were common to the family wide Annonaceae analysis (Figure 2 B) highlighting that the baiting kit can be used to simultaneously capture ingroup and outgroup taxa when targeting specific Annonaceae clades. This cross-family flexibility comes from the way we designed our baits. First, we used a top-down approach: baits were designed and optimized based on five species sampled from across the family. We thus identified orthologous exons that were common across our sample. Such approaches were already successful within other tropical plant families such as palms (Heyduk et al., 2016), where the baiting kit has been successfully used to sequence species in different clades of the family (Comer et al., 2016). In contrast, phylogenetic relationships within the large Neotropical genus *Inga* (Nicholls et al., 2015) or the mega diverse genus *Erica* (Kadlec et al., 2017) were reconstructed using baits designed from transcriptomes of species mostly within or closely related to target genera or study clade. In the latter studies, the baiting kit was shown to successfully cpature exons across the Ericaceae family too (Kadlec et al., 2017). Second, we also used a “degenerated” kit, meaning that for 282 (out of 469) identified orthologous exons we included different species-specific exon variants increasing capture capacity across the family. However, we did not undertake explicit analyses or resequencing to test if our degenerate kit improved capture across Annonaceae or not.

**Figure 5.**
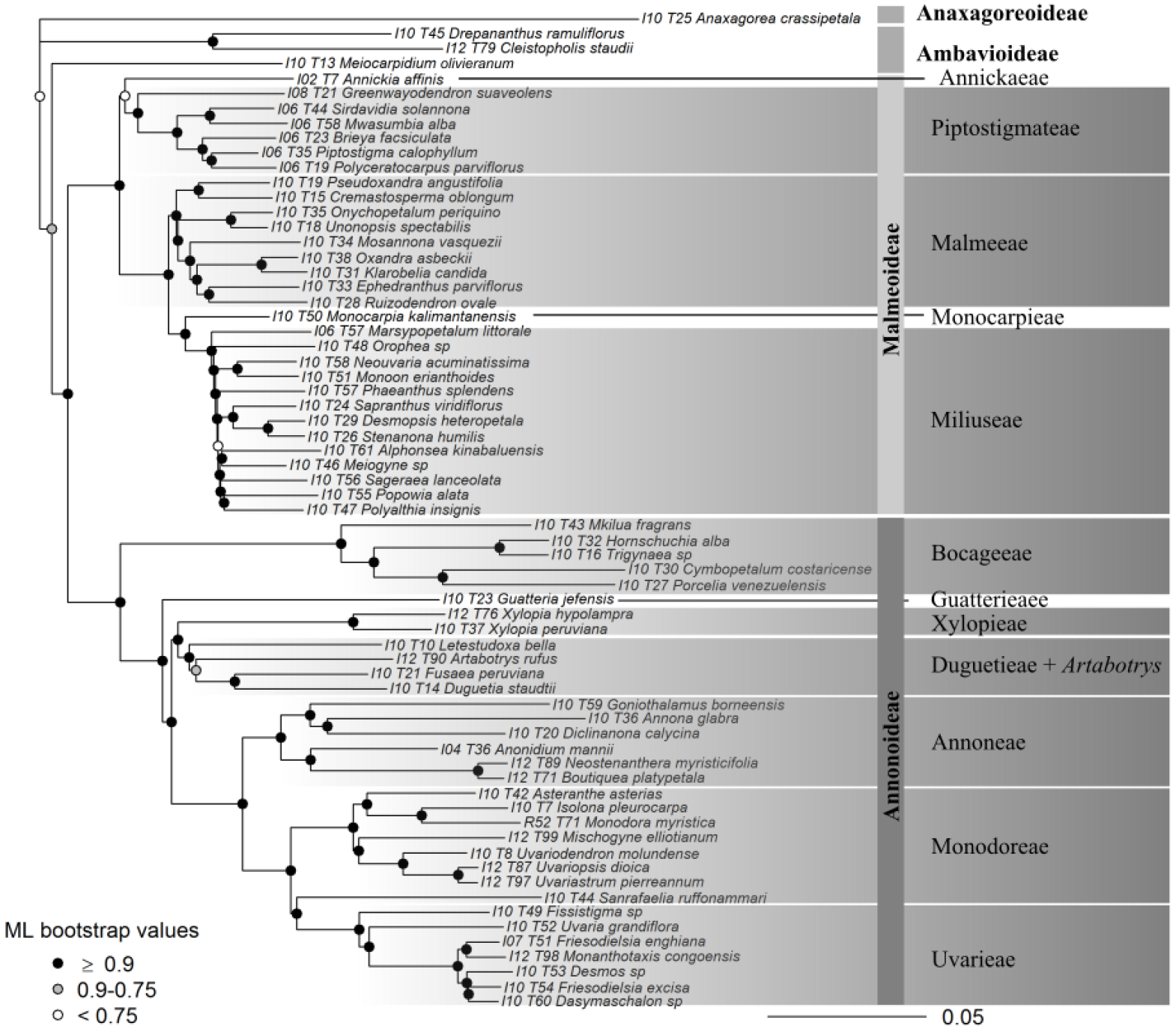
Maximum likelihood tree of Annonaceae based on 317 concatenated supercontigs. Supercontigs were concatenated after removal of putative paralogous loci to form a supermatrix to be used in RAxML. Grey scale colours at nodes depict branch support after 100 bootstrap replicates.

We recovered around 500-1000 bp of off-target sequence data (i.e. sequence data flanking our targeted regions). Off-target sequences are mostly composed of intronic data, but it has been shown that non-baited exon regions can also be captured in the process (Kadlec et al., 2017). Annonaceae do not have a sequenced genome yet, so it is hard to know precisely what we are recovering in off-target sequences. Recovery of off-target sequence data is an important property that allows the same baiting kit to be informative at several taxonomic levels, which is clearly shown by this and other studies (Kadlec et al., 2017). In a preliminary study we also used the same baiting kit to successfully capture 354 loci at infraspecific levels within *Greenwayodendron suaveolens* providing good phylogeographic resolution (unpublished).

**Figure 6.**
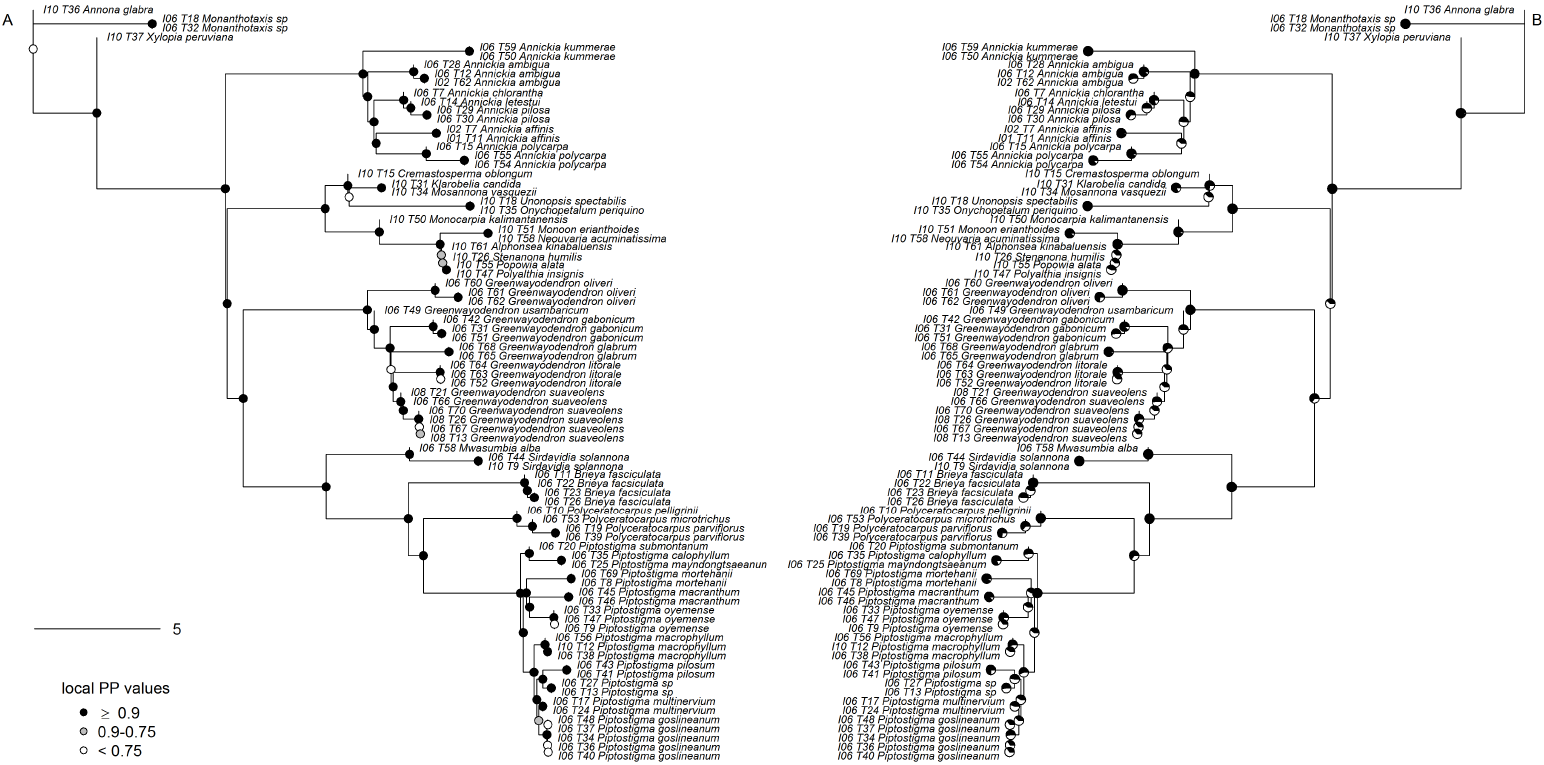
Species-level tree of Piptostigmateae constructed using ASTRAL. A. Tree inference was based on 356 supercontigs. Putative paralogous loci were identified and the entire locus was removed. Depicted on nodes are the local posterior probability (LPP) values. B. Identical tree to A but with quartet support represented on nodes. Black portion of pie charts represents the percentage of quartets in gene trees agreeing with this branch.

Finally, for two samples, DNA was extracted directly from herbarium specimens (INDEX6_TAG49 *Greenwayodendron usambaricum* and INDEX6_TAG50 *Annickia kummerae*) leading to good overall coverage and sequencing depth (Supplementary material). This confirms the utility of targeted enrichment approaches to sequence hundreds of nuclear genes for herbarium preserved material (Hart et al., 2016). DNA extracted from Annonaceae herbarium specimens have been successfully and routinely sequenced in the past even from 100+ year-old specimens. Thus our results are encouraging and suggest that our kit will be useful to tap into “genomic treasure trove” (Staats et al., 2013) of Annonaceae herbarium specimens.

### Substitution rates

Our Maximum Likelihood concatenated phylogenetic tree (Figure 5) based only on nuclear loci shows a clear difference in branch lengths between the two large Annonaceae subfamilies Annonoideae and Malmeoideae. Although no formal tests were undertaken here, our results suggest that this increased substitution rate within Annonoideae when compared to Malmeoideae affects the entire genome as already suggested by Hoekstra et al. (2017). This genome-wide among-lineage rate heterogeneity can have important impacts when estimating divergence times (e.g. Hoekstra et al., 2017; Bellot and Renner, 2014; Testo et al., 2018). This will need to be taken into account in further biogeographical analyses of the family (Hoekstra et al., 2017), for example using Random Local Clock approaches (Drummond and Suchard, 2010; Bellot and Renner, 2014; Testo et al., 2018).

**Figure 7.**
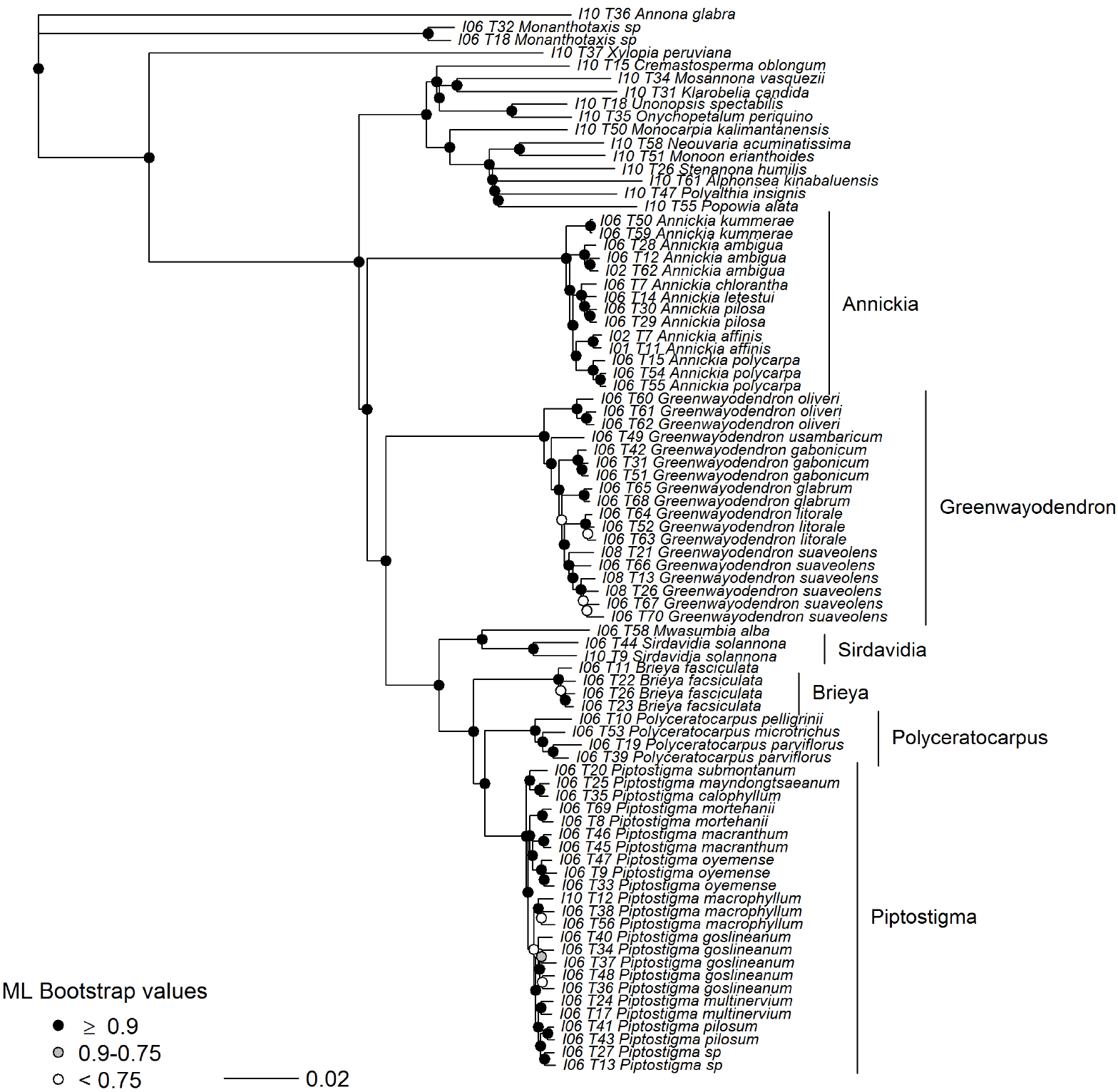
Maximum likelihood tree of Annonaceae based on 356 concatenated supercontigs. Supercontigs were concatenated after removal of individuals with putative paralogous sequences from alignments to form a supermatrix to be used in RAxML. Node colours show support based on 100 bootstrap replicates.

### Next generation Annonaceae phylogenetics

We present the first phylogenomic analyses of Annonaceae based on 317 (excluding paralogues) nuclear loci sampled for 65 out of 109 (60%) currently recognized genera (Figures 4, 5). Overall our results mostly agree with previous Annonaceae wide phylogenetic analyses solely based on plastid regions (Chatrou et al., 2012b; Guo et al., 2017a; Richardson et al., 2004; Su et al., 2008; Couvreur et al., 2011a; Guo et al., 2017b). Deep level relationships are congruent between the concatenated and gene tree approaches and all four sub families are recovered with maximum support (Figures 5, 4). The clade most affected by our methods is that of the genera of the Miliuseae tribe. Indeed, some of these intergeneric relationships are well supported in the concatenated approach, but not in the coalescent approach. Our results, however, do provide some new insights into the phylogeny of Annonaceae, which we describe here.

### Ambavioideae

#### Meiocarpidium

The position of the monotypic genus *Meiocarpidium* is moderately supported as sister to Annonaceae excluding *Anaxagorea* and Ambavoideae in the concatenated maximum likelihood tree (Figure 5), and weakly supported as sister to the rest of the Ambavoideae in the gene tree (Figure 4). About 1/3 of loci support the quartet relationship of *Meiocarpidium* as sister to the rest of the Ambavioideae (Figure 4 B) leading however to low LPP support. *Meiocarpidium* was already suggested to be early diverging within Annonaceae especially based on floral and pollen characters (Le Thomas, 1980, 1981) and its placement was problematic in past phylogenetic analyses. Indeed, this genus was recovered as sister to the rest of Ambavioideae based on eight plastid loci (Chatrou et al., 2012b; Guo et al., 2017a) with moderate bootstrap support. Here, despite substantial increase in sequence data the position of *Meiocarpidium* remains uncertain.

#### Guatteria

Our phylogenomic analysis sheds light on one of the major, unresolved phylogenetic questions within the Annonoideae: the position of the tribe Guatterieae (Erkens et al., 2009). This tribe consists solely of the species-rich Neotropical genus *Guaterria* (Maas et al., 2015). So far family-wide chloroplast-based analyses were not able to determine the position of this tribe with certainty (Erkens et al., 2009; Couvreur et al., 2011a; Chatrou et al., 2012b; Guo et al., 2017b). Here, we show with high support that Guatterieae is sister to all Annonoideae except the tribe Bocageeae (Figures 5, 4). This result, however, is only based on a single *Guaterria* species, and thus needs to be confirmed based on a larger sample of this genus.

#### Artabotrys

The unexpected position of the paleotropical genus *Artabotrys* renders the tribe Xylopieae paraphyletic. The relationship between tribes Xylopieae and Duguetieae are strongly supported in both our gene tree and concatenated analyses (Figures 5, 4). However, *Artabotrys* is always recovered as nested with high support within the Duguetieae clade (sampled genera *Letestudoxa*, *Fusaea*, *Duguetia*). These results contrast with what was inferences based on plastid data where *Artabotrys* was systematically recovered as sister with strong support to the large pantropical genus *Xylopia* (Richardson et al., 2004; Chatrou et al., 2012a; Thomas et al., 2015; Guo et al., 2017b). This relationship, however, was not recovered by cladistic analyses of morphological data (Doyle and Le Thomas, 1994) or *rbcL* sequence alone (Doyle et al., 2000). The tribe Xylopieae was erected following the phylogenetic analyses of eight plastid markers, but it was stressed that clear morphological synapomorphies between the two constituents genera were lacking (Chatrou et al., 2012b). Indeed, *Artabotrys* is a genus of lianas that has evolved special hook-shaped terminal inflorescences whereas *Xylopia* contains large tree species with non-hook shaped axillary inflorescences. It is beyond the scope of this paper to detail differences and similarities, but our results suggest that the phylogenetic placement of *Artabotrys* should be more thoroughly investigated, and the taxonomic implications reviewed. Increased sampling of both genera (5 out of 7 genera sampled here) and species could potentially help resolve this question.

#### Sanrafaelia

Our results suggest that the Tanzanian endemic genus *Sanrafaelia* is excluded from tribe Monodoreae, and is in fact sister with strong to the mainly climbing tribe Uvarieae (Figures 5, 4). Based on plastid markers *Sanrafaelia* together with its East African sister genus *Ophrypetalum* were recovered with moderate support as sister to a clade of African genera (Couvreur et al., 2008; Chatrou et al., 2012b; Guo et al., 2017b). Even though one synapomorphy was identified to support the erection of the Monodoreae tribe (sessile monocarps (Couvreur et al., 2008; Chatrou et al., 2012b)) it appears that this tribe might have to be revisited in the light of nuclear data, and especially the inclusion of the genus *Ophrypetalum.*

### Malmeoideae

#### Annickia

Our data suggest that the African tribe Piptostigmateae is potentially not monophyletic. Indeed, *Annickia* is resolved as sister with high support to the rest of the Malmeoideae in the gene tree analysis (Figure 4), but weakly supported as sister to the rest of the Piptostigmateae genera in the concatenated analysis (Figure 5). These results were also confirmed when we increased sampling in the tribe (Figures 7, 6) highlighting discordance between the gene tree and the concatenated analyses. In the gene tree analysis, not all loci recovered the *Annickia* sister to Malmeoideae relationship (41% in the Annonaceae analysis and 64% in the Piptostigmateae analysis). This clearly underlines some conflict in the proper placement of the genus. Nevertheless, this still translated into high LPP support for this branch because the alternative quartet topologies are recovered by significantly fewer loci. Substantial gene tree conflict cannot be accounted for by the concatenation approach (Liu et al., 2015) and this could be the reason why we recovered different relationships in the RAxML trees (Figures 5, 7). Discordance in phylogenetic relationships between concatenated and gene tree coalescent approaches based on phylogenomic data have been reported (Xi et al., 2014). We thus suggest that the gene tree analysis using ASTRAL provides a better inference of true relationships than the concatenated approach (Liu et al., 2015).

The placement of *Annickia* has been problematic in previous analyses as well. It has either been weakly inferred as sister to the rest of Malmeoideae (Doyle et al., 2000) or as sister to the rest of the Piptostigmateae genera based on several plastid loci (Pirie et al., 2006; Chatrou et al., 2012b; Guo et al., 2017b). Morphological and pollen character-based cladistic analyses in contrast underlined affinities of *Annickia* with other members of the Malmeoideae such as *Unonopsis* (Doyle and Le Thomas, 1994). *Annickia* was included in the tribe Piptostigmateae (Chatrou et al., 2012b) because it was resolved (but not supported) as sister to that tribe and probably because *Annickia* is, as all other Piptostigmateae genera, strictly African. *Annickia* presents several morphological differences with other Piptostigmateae genera such as in pollen (columelar versus granular infratectum) (Doyle and Le Thomas, 2012) and floral characters (one versus several ovules, more than 20 carpels versus less than 20) (Doyle and Le Thomas, 1994; Couvreur et al., 2009, 2015; Versteegh and Sosef, 2007). Based on the present phylogenomic analyses taking into account gene trees, both at generic and species level sampling, and morphology we suggest that the genus *Annickia* does not belong in the Piptostigmateae, and propose the description of a new tribe, Annickieae.

**Annickieae** Couvreur, **tribus nov**. - TYPE GENUS: *Annickia* Setten & Maas in Taxon 39(4): 678, 681 (1990). Trees, indumentum with single, bifid, trifid, fasciculate or stellate hairs; flowers bisexual, solitary, terminal; perianth actinomorphic, 6 free tepals in two opposite whorls of 3, differentiated in an outer whorl of 3 sepals and an inner whorl of 3 valvate petals. Stamens 110-175, filaments short, anthers extrorse; carpels 35-70, with a single ovule per carpel; monocarps free, stipitate. Comprising a single genus, *Annickia,* with eight species endemic to continental Africa (Versteegh and Sosef, 2007). The sister relationship of Annickieae with the rest of the Malmeoideae based on our gene tree analysis will have implications in understanding the evolution and biogeography of Annonaceae, especially within the Malmeoideae subfamily (e.g. Couvreur et al., 2011a; Thomas et al., 2017; Doyle and Le Thomas, 2012).

#### Miliuseae

Our results show that resolving the backbone of the Miliuseae tribe will be challenging even when using a phylogenomic approach. As when using plastid sequence data, branches leading up to the different sampled genera are short. However, most of the nodes are strongly supported in the concatenated approach (Figures 5) but the gene tree analysis shows high levels of discordance between loci (4). Nevertheless, our nuclear phylogenetic tree agrees with previous plastid-based analyses (Mols et al., 2004; Chatrou et al., 2012b; Chaowasku et al., 2014; Xue et al., 2012; Guo et al., 2017b). For example, *Sapranthus*, *Desmopsis* and *Stenanona* form a well supported clade which is also the case when using plastid markers (Guo et al., 2017b; Chatrou et al., 2012b; Ortiz-Rodriguez et al., 2016). The coalescent analysis highlights that most backbone relationships are supported by just one third of loci with low LPP support (Figure 4). Our results support the idea that the Miliuseae underwent a period of rapid diversification leading to widespread incomplete lineage sorting. However, it must be noted that our sampling of this tribe is far from complete, and more genera and species will need to be added to gain a true understanding of this clade.

### Piptostigmateae species-level phylogenetic tree

Besides the potision of *Annickia* (see above), our phylogenomic analysis of tribe Piptostigmateae (Figures 7, 6) recovers maximally supported relationships between genera, confirming previous plastid phylogenetic trees (Couvreur et al., 2009, 2015; Ghogue et al., 2017; Guo et al., 2017b). Our results confirm the generic status of *Brieya* as a phylogenetically separate genus from *Piptostigma* (Ghogue et al., 2017). The coalescent analysis (Figure 6) shows that the monophyly of all genera is supported by a high percentage of loci (>94%).

Relationships between species within genera are overall well supported in the concatenated analysis and with high LPP in the coalescent approach. Our results support the latest taxonomic revision of the genus *Greenwayodendron* with five phylogenetically different species (Lissambou et al. 2018, in press) instead of the long held view of two species (Le Thomas, 1969). This includes the recognition of the East African taxon *Greenwayodendron usambaricum* (Verdc.) Lissambou, Hardy & Couvreur as a distinct species from *Greenwayodendron suaveolens* (Engl. & Diels) Verdc‥ In addition, our results are also concordant with the latest revision of the genus *Piptostigma* (Ghogue et al., 2017). However, we did identify two individuals which appear to belong to a new undescribed species (labelled as sp. in figures 7, 6). Finally, using are baiting kit, all species were recovered as monophyletic. This suggests that overall, species circumscription in this tribe is well established.

## Conclusion

We show here that even within a tropical plant family with overall no available genomic resources, it is possible to generate nuclear markers useful for family wide and species level phylogenetic analyses. Thus, for other tropical plant lineages, having access to fresh material of a few non related species covering the diversity of the studied group is potentially enough to generate a phylogenetically useful nuclear baiting kit. In addition, these family specific kits could be used in tandem with the angiosperm universal baiting kit recently developed (Johnson et al., 2018). In the era of phylogenomics, it will be important not only to generate clade specific baits, but also try and include baits used across angiosperms for enabling easier sharing and compilation of data around the plant Tree of Life (Eiserhardt et al., 2018).

The nuclear baiting kit designed here enables the potential capture of up to 469 loci across Annonaceae. We show that over 170 loci can be captured across the whole family with good coverage. The kit is thus a good tool to generate a well resolved species-level phylogenetic tree of all Annonaceae species. This in turn, can be used to address numerous taxonomical or evolutionary questions within the family. It remains however to be tested if we can capture exons from closely related plant families such as Eupomataceae or other Magnoliales families.

The phylogenetic relations within the Miliusieae tribe remains hard to resolve even when using a large number of nuclear loci. This remains a problem for Annonaceae phylogenetics. Potentially, the addition of full chloroplast sequences or plastomes might provide some added resolution. Finally, it remains to be seen how much resolution and support we recover when reconstructing the phylogenetic relationships of species-rich genera such as *Guatteria* (Erkens et al., 2007; Maas et al., 2015) or *Goniothalamus* (Tang et al., 2015).

## Acknowledgments

This study was supported by grants from the Agence Nationale de la Recherche (AFRODYN: ANR-15- CE02-0002-01 to TLPC; MAGNIPHY: ANR-12-JVS7-0015-01 to HS); RHJE was supported via the Innovational Research Incentives Scheme (VENI, nr. 863.09.017; NWO-ALW, The Netherlands). We are grateful to Etienne Delannoy who coordinated the sequencing of the *Marsypopetalum* transcriptome.

